# Identification and Characterization of a Residual Host Cell Protein Hexosaminidase B Associated with N-glycan Degradation During the Stability Study of a Therapeutic Recombinant Monoclonal Antibody Product

**DOI:** 10.1101/2020.09.17.302679

**Authors:** Xuanwen Li, Yan An, Jing Liao, Li Xiao, Michael Swanson, Kirby Martinez-Fonts, Jorge Alexander Pavon, Edward C. Sherer, Vibha Jawa, Xinliu Gao, Simon Letarte, Douglas D. Richardson

## Abstract

Host cell proteins (HCPs) are process-related impurities derived from host organisms, which need to be controlled to ensure adequate product quality and safety. In this study, product quality attributes were tracked for several mAbs under the intended storage and accelerated stability conditions. One product quality attribute not expected to be stability indicating is the N-glycan heterogeneity profile. However, significant N-glycan degradation was observed for one mAbs under accelerated and stressed stability conditions. The root cause for this instability was attributed to Hexosaminidase B (HEXB), an enzyme known to remove terminal N-acetylglucosamine (GlcNAc). HEXB was identified by liquid chromatography–mass spectrometry (LC-MS) based proteomics approach to be enriched in the impacted stability batches from mAb-1. Subsequently, enzymatic and targeted multiple reaction monitoring (MRM) MS assays were developed to support process and product characterization. A potential interaction between HEXB and mAb-1 was initially observed from the analysis of process intermediates by proteomics among several mAbs and later supported by computational modeling. An improved bioprocess was developed to significantly reduce HEXB levels in the final drug substance. A risk assessment was conducted by evaluating the *in silico* immunogenicity risk and the impact on product quality. To the best of our knowledge, HEXB is the first residual HCP reported to have impact on the glycan profile of a formulated drug product. The combination of different analytical tools, mass spectrometry and computational modeling provide a general strategy on how to study residual HCP for biotherapeutics development.

## Introduction

Host cell proteins (HCPs) are process-related impurities derived from host organisms (e.g. Chinese Hamster Ovary cell (CHO), E. coli). Residual HCPs have potential adverse effects for both safety and efficacy of biologics,^1–2^ including induction of immunogenicity, acute toxicity, and impact on product and excipient stability. The regulatory expectations are residual HCPs have to be tested on a routine basis with a sensitive assay, and the limit of residual HCP needs to be controlled at release by methods that provide sufficient coverage. ^1–2^ Several high-risk HCPs have been reported to impact immunogenicity and product or excipient stability. Hamster phospholipase B-like 2 (PLBL2), is a common and problematic residual HCP found in CHO-derived monoclonal antibodies (mAbs); It was found to induce immunogenicity in patients at levels as low as 0.2-0.4 ppm, although no clinical adverse effect was detected in patients.^3^ In addition to immunogenicity concerns, lipases or enzymes with esterase like activity, like group XV lysosomal phospholipase A 2 isomer X1 (LPLA2) and lipoprotein lipase (LPL) have been reported to degrade formulation excipients such as polysorbate 80 or 20.^4–5^ Trace levels of other residual HCPs, such as Cathepsin D ^6–7^ and Cathepsin L,^8^ were reported to have impact on drug product degradation by inducing fragmentation. The impact of a residual HCP on protein modifications of biologics in real-time stability studies has not been reported before.

Glycosylation is a critical posttranslational modification (PTM) in biologics that affects several relevant functions, such as pharmacokinetics, pharmacodynamics, and immunogenicity.^9–11^ For instance, sialic acid is present on N-linked or O-linked glycans. Both variants can extend biologics half-life, while high mannose glycosylation is known to reduce half-life.^12^ Furthermore, in cancer immunotherapy, the absence of core fucose on some IgG1 mAbs has been shown to increase fragment crystallizable (Fc) binding affinity to its receptor, leading to enhanced antibody-dependent cellular cytotoxicity (ADCC).^13^ Likewise, nonhuman and nonmammalian glycans, such as galactose-α1,3-galactose (α-Gal) and high-mannose N-glycans, can induce immunogenicity response.^9–10^ Therefore, it is critical to properly control glycosylation during a biologics manufacturing process.

Glycosylation profile is known to be impacted by multiple steps in biologics production, such as molecule sequence design, cell line engineering, upstream and downstream process.^9–10^ For example, the N-glycan processing enzymes of the host cell line can be modified by strain engineering to alter glycosyltransferases or glycosidases levels. It has been reported that reduction of non-human and potentially immunogenic glycans has been achieved by elimination of α-1,6-fucosyltransferase activity in CHO cells by removing α-1,3-galactosyltransferase and CMP-Neu5Ac hydroxylase from the strain.^14^ In another example, ADCC was enhanced through the introduction of beta-1,4-mannosyl-glycoprotein 4-beta-N-acetylglucosaminyltransferase, which prevented fucosylation and production of bisecting N-glycan.^14^ Additionally, cell culture conditions in upstream process, such as pH, temperature, CO_2_, and ammonium ion concentration, have been reported to have impact on glycosylation heterogeneity. In downstream process, major glycosylation isoforms may be separated according to charge variations with ion-exchanging chromatography (IEX) or based on hydrophobicity by hydrophobic interaction chromatography (HIC).^15^

Unit operations or various purification steps of the downstream process are designed to remove the majority of HCPs. The acceptance level of HCPs in the final drug product is typically in the range 1-100 ppm (1-100 ng/mg product) as measured by enzyme linked immunosorbent assay (ELISA).^2^ Due to assay limitation of ELISA, including antibody coverage, there may be still significant number of HCPs detected in final drug products which can have adverse effect on product quality or patient safety. Alternative analytical assays, such as liquid chromatography– mass spectrometry (LC-MS) based approaches, provide complementary characterization of HCPs for their composition and relative abundances.^1^

In this study, we are reporting a residual host cell enzyme, Hexosaminidase B (HEXB), as the cause of N-glycan GlcNAc degradation observed in mAb stability studies. The effect and level of HEXB in mAb-1 were tested with a β-N-Acetyl-glucosaminidase (NAG) enzymatic assay and LC-MS based MRM assay. The potential interactions responsible for co-purification were investigated through a molecular modeling approach. Additionally, we evaluated the epitope content and similarity of the protein to difficult-to-remove HCPs to understand the clinical risk of the co-eluted HEXB. In the end, an improved process was able to reduce this HCP level and mitigate its impact on glycan profile.

## Materials and Methods

### Materials

Dithiothreitol (DTT) and iodoacetamide (IAM) were purchased from Thermo Pierce (Rockford, IL). Tris-HCl buffer and high-performance liquid chromatography (HPLC) grade solvents, methanol, acetonitrile (ACN), formic acid (FA), were purchased from Thermo Fisher Scientific (Waltham, MA). The custom stable isotopic labeled (SIL) surrogate peptide (GILIDTSR, AQUA Ultimate grade) with 13C and 15N labeled arginine (R*), and non-labeled surrogate peptide were synthesized from Thermo Fisher Scientific (Waltham, MA). The mAbs are from the pipeline of Merck & Co., Inc., Kenilworth, NJ, USA. Sequencing grade modified trypsin was purchased from Promega (Madison, WI). Ammonium bicarbonate and urea were purchased from JT Baker (Center Valley, PA). Deep well 1-mL 96 well plate (Protein LoBind) was purchased from Eppendorf (Hauppauge, NY). Recombinant mouse HEXB protein (His Tag) was obtained from MyBioSource (San Diego, CA). Recombinant proteins alcohol dehydrogenase (ADH) from yeast, carbonic anhydrase ii (CA2) from *Bos Taurus*, myoglobin (MB) from equine heart, phosphorylase-b (PYGM) from rabbit muscle and enolase (ENL) from baker’s yeast, and enzyme activity assay kit for β-N-Acetyl-glucosaminidase (NAG) were purchased from Sigma (St. Louis, MO). Recombinant NAG enzyme was from New England Biolabs (Beverly, MA). Instant PC kit was from Agilent (Palo Alto, CA).

### N-glycan Quantification Assay

The quantification of N-glycan species was analyzed using an Instant PC kit according to the manufacturer protocol. The samples were prepared using a high-throughput 96 well plate-based method. N-linked glycans were hydrolyzed from the protein using N-Glycanase enzyme, which cleaved the bond between asparagine and β-N-acetylglucosamine. The free glycans were subsequently labeled with the dye Instant PC and purified with a cleanup column. The labeled glycans were analyzed using normal phase UPLC with fluorescence detection.

### Total HCP by ELISA

Total HCP content in mAb-1 DS from Process 1 was measured by a CHO HCP ELISA kit 3G (F550, Cygnus, Southport, NC) according to the manufacture protocol. The drug substance is diluted to approximately 5 mg/mL in commercial sample diluent. Diluted samples and standards were reacted in microtiter strips coated with an affinity purified capture antibody. A second horseradish peroxidase enzyme labeled anti-CHO-HCP antibody was reacted simultaneously resulting in the formation of a sandwich complex. After incubating for two hours, the wells were washed to remove any unbound reactants and a substrate containing tetramethyl benzidine (TMB) was added. The TMB yielded a color-change upon reaction. The amount of hydrolyzed substrate was directly proportional to the concentration of HCP present which was quantified using the standard curve.

### HCP Identification and Relative Quantification by Proteomics

Residual HCPs in process intermediates and drug substance of several mAbs were identified by LC-MS-based proteomics approach using limited enzyme digestion for HCP enrichment.^16^ Briefly, 1 mg of protein was mixed with 10 μL of Tris-HCl (1 M, pH 8), 2.5 μg of trypsin, 10 μL of stocking protein mixture and LC-MS grade water to a total of 200 μL. The 100× stocking protein mixture used for system suitability testing and relative quantification was made by mixing 5 non-CHO recombinant proteins. The final working concentration of each standard protein compared to DS is 200 ppm for ADH, 100 ppm for CA2, 50 ppm for MB, 25 ppm for PYGM and 10 ppm for ENL. Proteins were digested at 37 °C overnight, and reduced with 2 μL of 50 mg/mL DTT at 80 °C for 10 min. A large portion of undigested proteins was removed by centrifugation at 11,000 *x g* for 10 min at room temperature. The collected supernatant was mixed with 3 μL of 20% FA before LC-MS analysis.

LC-MS-based proteomics analysis was run on an ACQUITY UPLC H-Class system (Waters, Milford, MA) coupled with Q Exactive™ HF-X Hybrid Quadrupole-Orbitrap™ mass spectrometry (Thermo, San Jose, California). The column was ACQUITY UPLC Peptide CSH C18 column (130 Å, 1.7 μm, 1 × 150 mm, Waters, Milford, MA) with flow rate at 50 μl/min and column temperature at 50 °C. Mobile phase A was LC-MS grade water with 0.1% FA, and mobile phase B was LC-MS grade ACN with 0.1% FA. The gradient started with 1%B for 5 mins, and increased to 5% at 6 min, and changed to 26% at 85 mins. The column was washed with 90%B from 90 mins to 105 mins, followed by 2 cycles of zig-zag washing step from 5% B to 90% B to reduce carry-over peptides. The MS data was acquired in data dependent analysis (DDA) mode with MS1 scan range from 300 to 1800 m/z and top 20 for MS/MS fragmentation. The MS1 resolution was 60,000, AGC target 1e^6^, maximum IT for 60 ms. The MS2 resolution was 15,000, AGC target 1e^5^ and maximum IT for 100 ms, isolation window at 1.4 m/z and NCE at 27. The dynamic exclusion was set for 20 s. The ESI source was run with sheath gas flow rate at 35, aux gas flow rate at 10, spray voltage at 3.8 kV, capillary temperature at 275 °C, Funnel RF level at 35 and aux gas heater temperature at 100 °C.

MS raw data was searched again the internal CHO FASTA database from Merck & Co., Inc., Kenilworth, NJ, USA customized with mAb and the 5 recombinant protein sequences using Proteome Discoverer 2.2 (Thermo, San Jose, California). The precursor mass tolerance was set at 15 ppm and fragment mass tolerance at 0.02 Da. The dynamic modification was set for M oxidation and maximum 3 modification. The target FDR for peptide identification was 0.01 and protein identification filter required at least 2 unique peptide identification. The total MS1 peak area from all identified peptides were used for the estimation of protein relative abundance.

### NAG Enzymatic Activity

NAG activity was measured by the hydrolysis of NAG substrate 4-Nitrophenyl N-acetyl-β-D-glucosaminide (NP-GlcNAc) according to the protocol supplied with the kit. Briefly, 10 mg of NP-GlcNAc was dissolved and mixed in 10 ml of 0.09 M citrate buffer solution to make the reaction buffer. Various mAbs or the same mAb with different concentrations were mixed with reaction buffer and incubated at 37 °C for 30 mins. The reaction was stopped by adding stop solution supplied in the kit. This enzymatic activity was measured colorimetrically at 405 nm. NAG enzyme from *Canavalia ensiformis* (Jack bean) supplied in the kit was used as a control to calculate the absolute enzyme activity.

### NAG Enzyme Incubation

Various concentrations of recombinant NAG enzyme (5, 10, 20 and 40 U/mL) from *Streptococcus pneumoniae* were incubated with mAb-1 from Process 2 or mAb-2 at 37 °C for 1 hr. After the reaction, the glycan profiles on mAb-1 and mAb-2 were measured by the Instant PC glycan assay mentioned above.

### Absolute Quantification of HEXB by LC-MRM Approach

Process intermediate samples for mAb-1, including harvested cell culture fluid (HCCF), Protein A affinity chromatography-purified product (PAP), filtered neutralized viral inactivation product (FNVIP), second column-purified product, third column-purified product, and various DS from Process 1 and Process 2, were tested by MRM as we did for lipase quantification.^17^ The total protein concentrations of all the samples were determined by measuring the absorbance at 280 nm using a UV-Vis spectrometer. Samples were all diluted to 5 mg/mL with PBS buffer before sample preparation.

A standard curve was made using mAb-2 without detectable HEXB as a matrix for quantification. The mAb was diluted to 5 mg/mL with PBS buffer (pH 7.4) and spikes of various amounts were used to build the standard curve. A stock solution containing HEXB was spiked into the matrix at 1.00, 2.00, 4.00, 10.0, 40.0, 100, 200, 500, and 800 ng/mg of DS (ppm) for the standard curve. Briefly, 30 μL aliquots of each sample including standards and matrix blank samples were added to a 1-mL 96-well polypropylene plate. A 50 μL aliquot of freshly prepared denaturation and reduction solution (10 M urea and 8 mM DTT in 50 mM ammonium bicarbonate solution), was added to each well, mixed by vortex, and incubated for 1 hour at 37 °C in a Thermo-Mixer (Eppendorf, Hamburg, Germany) at 500 rpm. A 10 μL aliquot of 135 mM IAM was added to each well, mixed by vortex, and incubated for 1 hours at room temperature in dark. Freshly prepared trypsin digestion buffer (300 μL at 33 μg/mL) was then added to each well and incubated for 13 hr at 37°C. Following trypsin digestion, a 20 μL aliquot of SIL peptide solution (10 nM) as internal standard (IS) was added and a 10 μL aliquot of 50% FA in water was added to each well to quench the digestion.

The quantitative analysis was performed on an UPLC-MS/MS instrument which consisted of a Waters H-class UPLC system and a Waters TQS mass spectrometer (Waters, Milford, MA). Aliquots of digestion samples (10μL) were injected onto an Acquity UPLC BEH C18 column (1.7 μm, 2.1×50 mm, Waters, Milford, MA) held at 40 °C. Mobile phase A was water with 0.1% FA and mobile phase B was ACN with 0.1% FA. The following gradients were applied at a flow rate of 300 μL/min for a total run time of 9 min: an initial isocratic flow of 5% B for 1 min was followed by a linear gradient from 5% B to 35% B in 3 min, a wash step at 95% B for 1.8 min each, and then equilibration with 5% B for 1.8 min to starting conditions to prepare the column for the next injection. Electrospray ionization (ESI) source parameters were set at 2.5 kV for capillary voltage, 35 V for cone voltage, 50 V for source offset, 500 °C for desolvation temperature, 850 L/hr for desolvation gas flow, 150 L/hr for cone gas flow, and 0.2 mL/min for collision gas flow. Data was collected and processed using MassLynx and TargetLynx software (version 4.1). Calibration curve regression and study samples were performed on TargetLynx (version 4.1). The calibration curves (analyte peak area/IS peak area versus analyte concentration) were constructed using linear regression fit (y = ax + b), and a weighting factor of 1/x^2^ was applied to the regression.

### Homology Models of HEXB and Antibody Fab Domains and HEXB/Antibody Complex

The homology model of HEXB from CHO cells was constructed using the X-ray structure of human HEXB ^18^ (PDB id: 1NOU.pdb) as the template. Molecular Operating Environment (MOE) software from CCG (Chemical Computing Group ULC, Montreal, QC, Canada, 2019) was used for model building.

Homology models of Fab domain of antibodies were built using Antibody Modeler of MOE, where the templates were selected with the X-ray structures of the most homologous antibodies from a collection of several thousand well curated antibody structures in the PDB, including framework and CDR loops. Sequence alignment and model building followed the procedures outlined by MOE Antibody Modeler.

The MOE protein-protein docking program was utilized to generate proposed structural models of the HEXB/antibody complexes. While exploring possible binding interfaces between the antibody and HEXB, the binding surfaces of the antibody were restrained to the antibody CDRs.

### Binding Energy and Buried Surface Area

Complex structures of HEXB/antibody were prepared for MMGBSA binding energy calculations using Maestro (Schrödinger 2020-2 release, 23, Schrödinger, LLC, New York, NY, 2020). The atomic coordinates of each model were imported and refined using the Protein Preparation Wizard with default parameters and settings. The protonation states were determined and set for all titratable residues at pH 7.0. Next, a restrained MacroModel energy minimization of all atoms was performed prior to Prime all atom minimization. Finally, Prime MMGBSA binding free energy was computed for each minimized model. The buried surface area upon binding of each complex was computed using Maestro.

### *In Silico* Immunogenicity Evaluation of HEXB in Patients

The foreign epitope content associated with HEXB and other HCPs was analyzed using the EpiVax’s ISPRI_HCP program. The output was in the form of an EpiMatrix score which is based on the number of epitopes, their predicted binding affinity for prevalent HLA-DR alleles, and the size of the protein. Predicted binding affinity is based on a Z-score calculation which is a measure of a 9-mer peptide’s predicted binding to an HLA-DR allele pocket. The top 5% binders have a Z-score ≥ 1.64. Epitopes were compared to a corresponding human homolog sequence to determine if the epitopes were present in humans or unique to the CHO cells.

Identified epitopes with binding potential were further analyzed for their predicted T cell receptor (TCR) facing residues. TCR facing residues were analyzed for conservation with human proteins in the JanusMatrix tool from EpiVax. A lower Janus score indicates lack of cross-reactive epitopes and a higher risk of immunogenicity.

## Results

### N-glycan Degradation of mAb1 in a Stability Study

N-glycan degradation was observed for an IgG1 molecule, mAb-1 during a real-time stability study at selected time points and temperatures. Significant degradation of GlcNAc in G0F and G1F and increase of G0F-GlcNAc and Core F because of GlcNAc removal were observed after 6 months of storage at 40°C by an Instant PC glycan assay (Figure 1 and Table 1). There was minor change at 6 months and 25°C, and no significant change of N-glycan profile was observed up to 12 months and 5°C (Supplementary Table 1 and Table 2). The glycan profile change was also detected by intact mass analysis by LC-MS (data not shown). In contrast, there was no major change of N-glycan in the stability study of mAb-2, mAb-3 and mAb-4. The N-glycan profile of mAb-1 stayed similar following force degradation conditions including pH and light stress (data not shown), suggesting this N-glycan stability is likely not related to chemical stress. Therefore, it was proposed that residual HCP was responsible for the change in N-glycan profile in mAb-1. The process development of mAb-1 was improved by incorporating a second polishing column to further remove HCPs. Following this process change, there was no significant change in the N-glycan profile in similar stability studies (data not shown).

**Figure 1.**
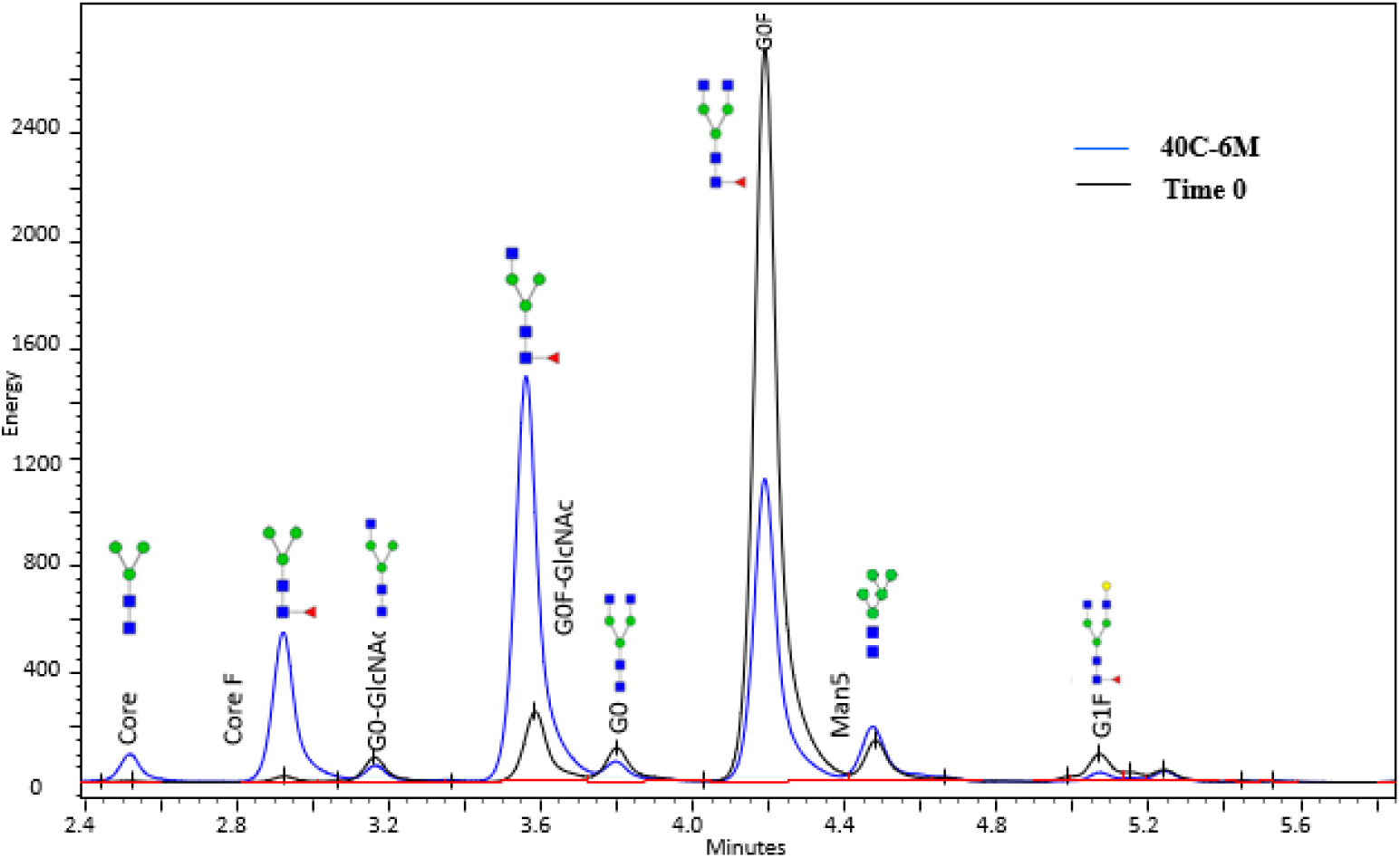
N-glycan profile change during stability study. The N-glycan trace from mAb1 between time 0 and 6 months at 40°C were presented.

**Table 1.** Percent peak area for N-glycan profile of mAb-1 stability sample at 40 °C measured by Instant PC assay.

|  | G0F | G0F-GlcNAc | CoreF | Core | G0 | G1F | Man5 | G0-GlcNAc |
| --- | --- | --- | --- | --- | --- | --- | --- | --- |
| 40°C-T0 | 75.1 | 7.1 | 0.8 | 0.2 | 3.5 | 3.5 | 4.1 | 2.3 |
| 40°C-3M | 47.0 | 27.4 | 10.1 | 2.1 | 2.7 | 1.5 | 4.1 | 1.1 |
| 40°C-6M | 30.4 | 39.4 | 13.1 | 2.3 | 2.1 | 1.1 | 4.3 | 1.8 |

### Identification and Relative Quantification of Hexosaminidase B for N-glycan Change

The residual total HCP level by a CHO ELISA kit for mAb1 DS from Process 1 ranged from 18 to 29 ppm (n=3). To identify which residual HCPs were the root cause for N-glycan GlcNAc removal, mAb-1 DS from Process 1 was analyzed by an LC-MS based proteomics approach. HEXB was the only glycosidase identified, with 24 unique peptides (p < 0.01) and sequence coverage at 47.9% (Figure 2). To determine the relative abundance of HEXB by proteomics, 5 non-CHO recombinant proteins ranging from 10 ppm to 200 ppm were spiked in each sample to make the standard curves. The correlations (R^2^) of the MS1 peak areas and the amount of the 5 spike-in proteins are over 0.95 (Figure 3). Relative quantification of HEXB calculated from the standard curve was determined to be 658 ppm from two biological replicates (Table 2). This residual HCP was not identified in the drug substances of mAb-2, mAb-3 and mAb-4. After process optimization to remove HCPs in mAb-1 from Process 2, HEXB could still be identified by proteomics, but coverage was reduced to only 6 unique peptides identified. The relative quantification from proteomics analysis is calculated as 16 ppm from the standard curve (Table 2).

**Figure 2.**
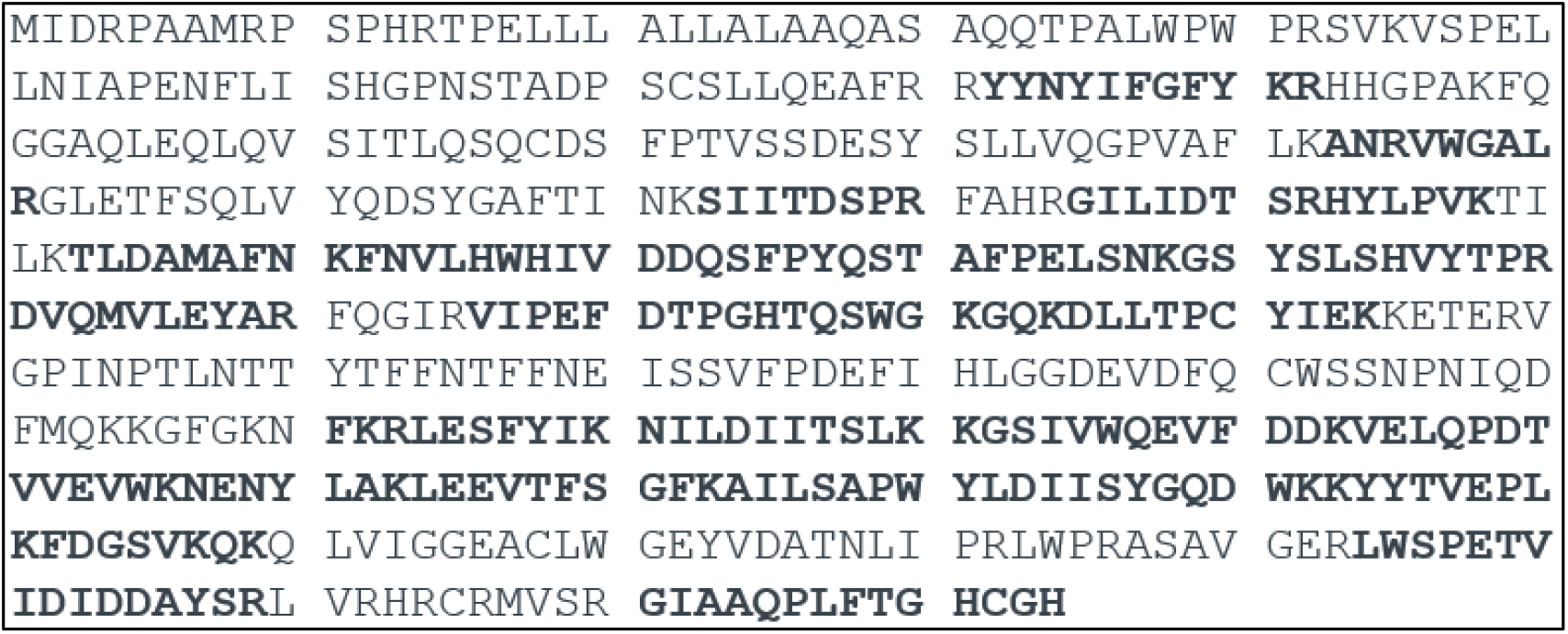
HEXB identification by LC-MS based proteomics approach. The amino acid of identified peptides by MS are in bold.

**Figure 3.**
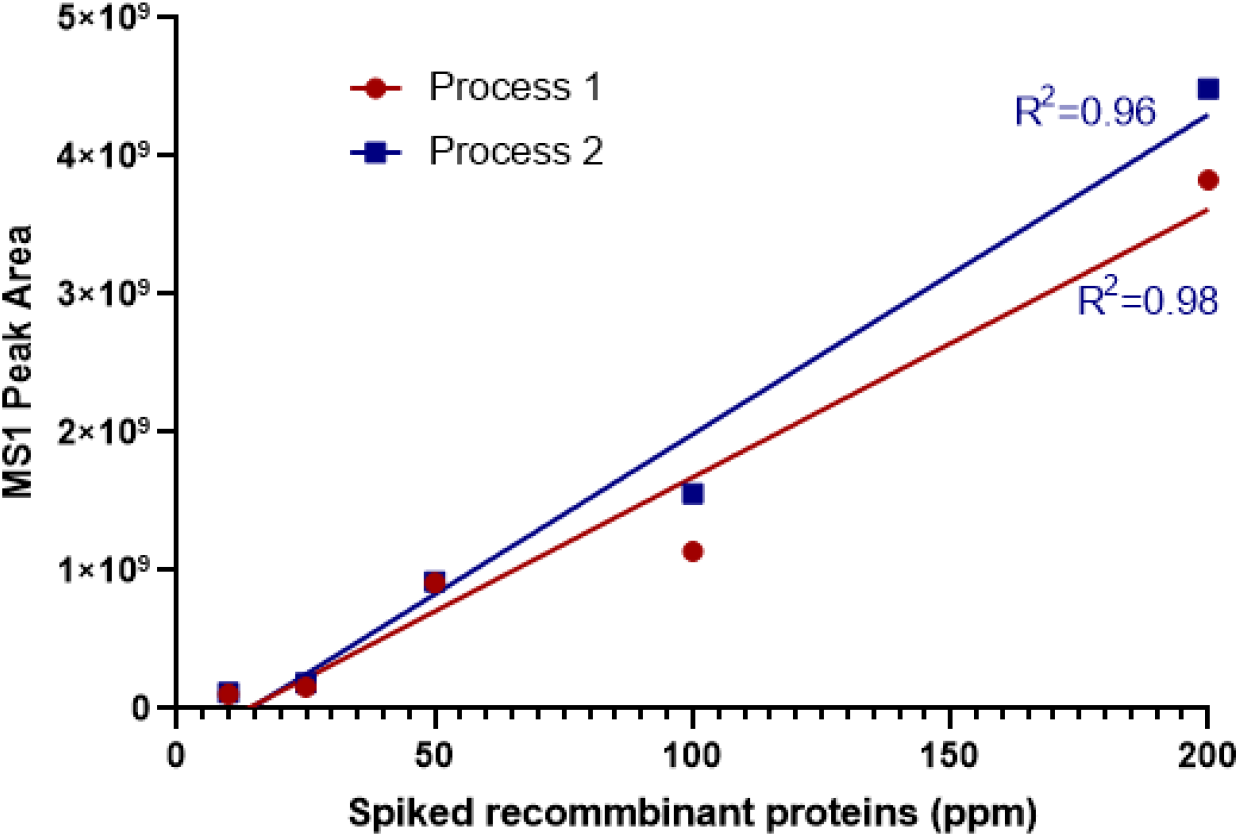
Standard Curves for HCP quantification from LC-MS based proteomics study. Five non-CHO recombinant proteins ranging from 10 to 200 ppm were spiked in mAb-1 from Process 1 and Process 2 before digestion. The extracted MS1 peak area of each protein was plotted with the spike-in amount in ppm.

**Table 2.** The relative quantification of HEXB by LC-MS based proteomics approach.

|  | Process 1 |  |  | Process 2 |  |  |
| --- | --- | --- | --- | --- | --- | --- |
|  | Replicate 1 | Replicate 2 | Average | Replicate 1 | Replicate 2 | Average |
| MS1 Peak Area | 1.10E+10 | 3.73E+09 | 7.35E+09 | 7.38E+07 | 2.85E+07 | 5.12E+07 |
| Relative Abundance (ppm) | 565 | 752 | 658 | 20 | 11 | 16 |

### Measurement of NAG Enzyme Activity

Hexosaminidase (EC 3.2.1.52), also called N-acetyl-β-D-glucosaminidase (NAG) in other species, have been reported to catalyze the hydrolysis of terminal GlcNAc residues from a variety of substrates. To determine the role of HEXB in mAb-1 glycan change, an enzyme activity assay was developed to measure HEXB activity on GlcNAc hydrolysis in mAb-1 from Process-1 and Process-2 using NP-GlcNAc as the substrates. As shown in Figure 4, NAG hydrolysis activity is highly correlated with mAb-1 concentrations from both processes. mAb1 from Process 1 has much higher NAG activity than that from Process 2. As expected in Table 3, mAb-1 from Process-1 had the highest NAG activity among the 4 mAbs. mAb-1 from Process 2 showed more than 20-fold decrease in NAG activity. The decrease in NAG enzyme activity is consistent with the decreases in HEXB concentration in Process 2. NAG enzyme activities in mAb2, mAb3 and mAb4 are extremely low as expected from their below limit of identification of HEXB by proteomics.

**Figure 4.**
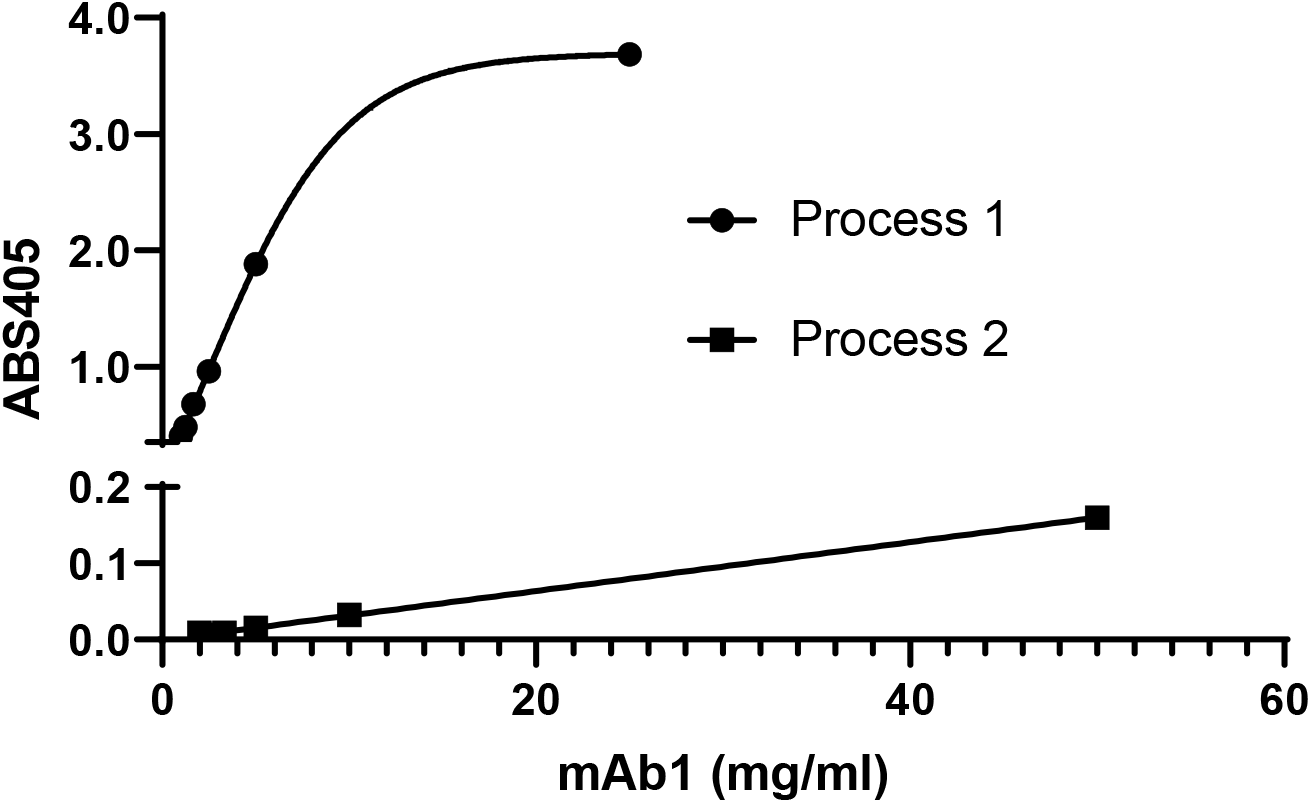
Enzymatic assay to measure NAG Enzyme Activity in mAb-1 from Process 1 and Process 2. Various amount of mAb-1 from Process 1 and Process 2 were incubated with NAG substrate NP-GlcNAc at 37 °C for 30 mins. This enzymatic activity was measured colorimetrically at 405 nm (n=3).

**Table 3.** NAG activity and relative quantification of HEXB by proteomics approach among 4 mAbs.

| DS | Activity (unit/ml) | HEXB (ppm) |
| --- | --- | --- |
| mAb-1 process 1 | 1.556 | 658 |
| mAb-1 process 2 | 0.068 | 16 |
| mAb-2 | 0.003 | ND |
| mAb-3 | 0.004 | ND |
| mAb-4 | 0.009 | ND |

ND: Not Identified

### Confirmation of NAG Like Activity as the Root Cause for GlcNAc Removal in mAbs

Due to the lack of high-quality recombinant CHO HEXB protein, a NAG enzyme with the same specificity as HEXB was used to confirm NAG enzyme activity as the root cause of N-glycan GlcNAc removal. Different concentrations of NAG were spiked in mAb-1 from Process 2 and in mAb-3. As shown in Figure 5A and 5B, the GlcNAc removal on mAbs can be replicated by adding NAG enzyme measured by the Instant PC N-glycan assay. Incubate of NAG at 40°C as short as 1 hour resulted in concentration-dependent decrease of G0F and increase of G0F-GlcNAc and CoreF, consistent with terminal GlcNAc removal from both mAbs.

**Figure 5.**
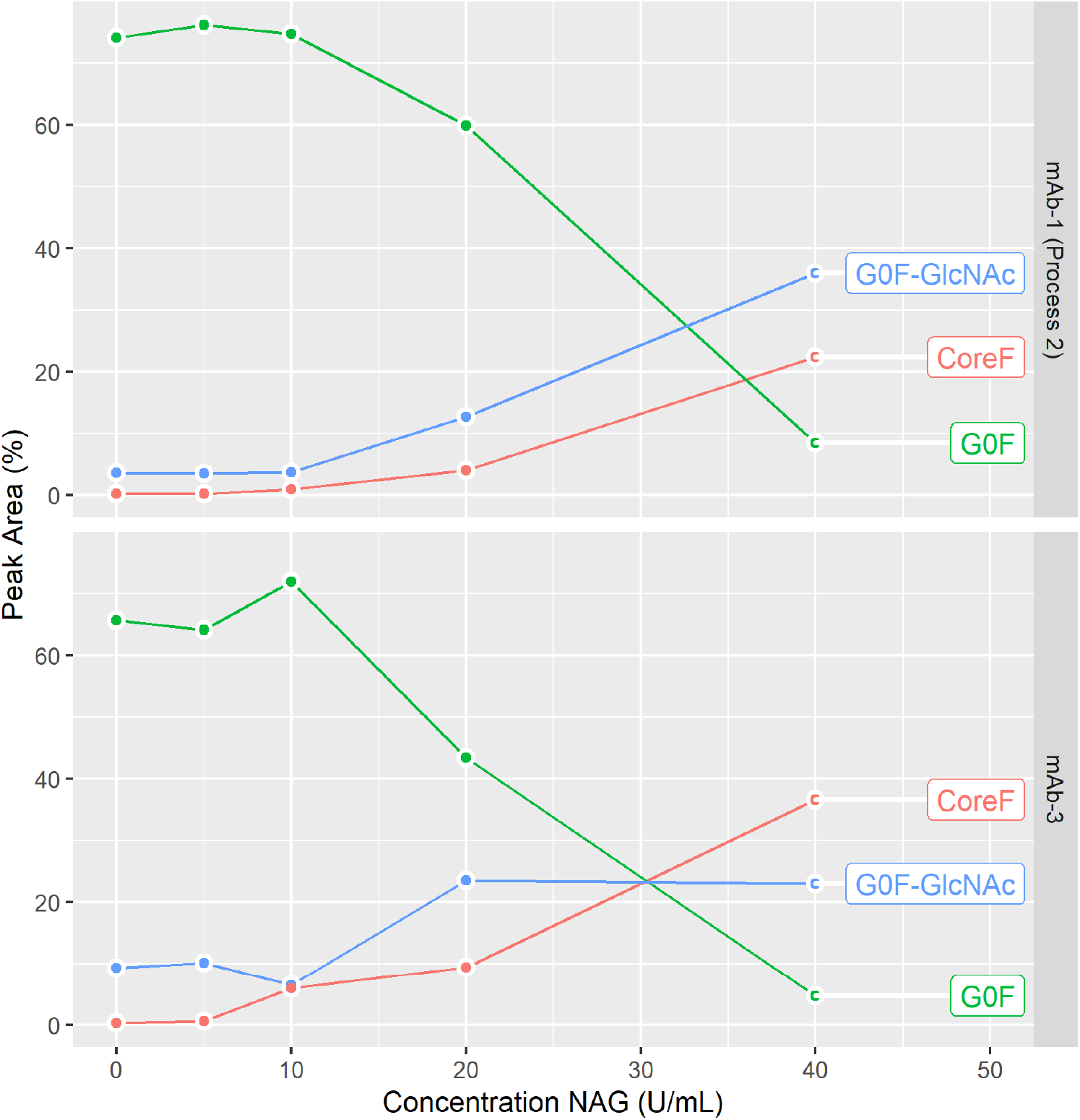
NAG incubation with mAb-1 from Process 2 or mAb-3. The NAG incubation with various concentrations from 5 U/mL to 40 U/mL was performed at 40°C for 1hr. N-glycan profile was measured by an Instant PC method. % of peak area is calculated.

### Absolute Quantification of HEXB by LC-MRM Approach

To help increase process understanding around the clearance of HEXB in mAb-1, an isotope dilution LC-MRM based method was developed to measure the absolute level of HEXB in process intermediates from Process 1 and Process 2. As shown in Table 4, HEXB is decreased along with the purification steps. HEXB levels in the final DS of Process 1 and Process 2 were determined to be 214.9 ppm and 3.8 ppm, respectively. The abundance change is consistent with the relative quantification from the proteomics analysis and the enzymatic activity.

**Table 4.** HEXB quantification (ppm) determined by LC-MS based MRM approaches.

| Samples | MRM (ppm) |
| --- | --- |
| HCCF | 794.4 |
| PAP | 255.1 |
| FNVIP | 53.6 |
| Column 1 | 17.4 |
| Column 2 | 0.4 |
| DS1 (Process 2) | 3.8 |
| DS2 (Process 2) | 3.8 |
| DS1 (Process 1) | 214.9 |
| DS2 (Process 1) | 219.7 |
| DS3 (Process 1) | 213.9 |

### HEXB Abundances in Process Intermediates from mAbs

To gain additional understanding as to why HEXB was specifically observed in mAb-1 as a residual HCP, but not in mAb-2, mAb-3 and mAb-4, process intermediates from those 4 mAbs were tested by a proteomics approach. Based on proteomics data from the 4 mAbs (Table 5) and other mAb molecules in the pipeline (data not shown), there are no major differences of HEXB levels in the harvest cell culture fluid (HCCF) for mAbs from different cell lines. Surprisingly, the HEXB level was significantly higher in the Pro-A product (PAP) from mAb-1 compared to other mAbs. HEXB was not identified in mAb-2 and mAb-4 in PAP, and only a very low level of HEXB was identified from mAb-3 in PAP. As observed earlier in the proteomics analysis, HEXB was only identified in the mAb-1 DS, but not in other mAb DS lots.

**Table 5.** Potential interaction between mAb-1 and HEXB during purification revealed from proteomics analysis.

| Molecule | mAb-1 | mAb-2 | mAb-3 | mAb-4 |
| --- | --- | --- | --- | --- |
| IgG isotype | IgG1 | IgG1 | IgG4 | IgG4 |
| Cell line | 1 | 1 | 2 | 2 |
| HCCF | 4.85E+08 | NT | 1.56E+08 | NT |
| PAP | 7.34E+09 | ND | 9.75E+06 | ND |
| DS | 1.53E+07 | ND | ND | ND |

NT: not tested; ND: not detected

### Modeling of HEXB Interaction with mAbs

To investigate the potential interactions of HEXB with mAb-1, computational modeling was conducted to understand the possible interactions at a molecular level. First, the homology model of HEXB from CHO cell was constructed using the human HEXB X-ray structure as the template. The two proteins are highly homologous with a sequence identity of 73%. Multiple complex structures generated using MOE protein-protein docking with the homology models of HEXB and antibody Fab domain were ranked using the MOE docking score and clustered based on potential epitopes on HEXB. Since the binding interface was restrained to antibody CDRs, the docking poses display interactions mostly from residues in CDRs. The proposed complex structure of HEXB/mAb-1 was selected among different docking poses based on top docking score. This complex model (Figure 6A) suggests that the binding epitope of HEXB for mAb-1 encompasses domain II, near the single disulfide bond in domain II.^18^ Furthermore, the epitope is away from the HEXB dimer interface to allow HEXB dimer formation, a necessary determinant of its enzymatic activity. The binding models of HEXB with the other three antibodies, mAb-2, mAb-3 and mAb-4 are shown in Figure 6B, 6C and 6D, where the binding interfaces occupy a similar region on HEXB as in mAb-1.

**Figure 6.**
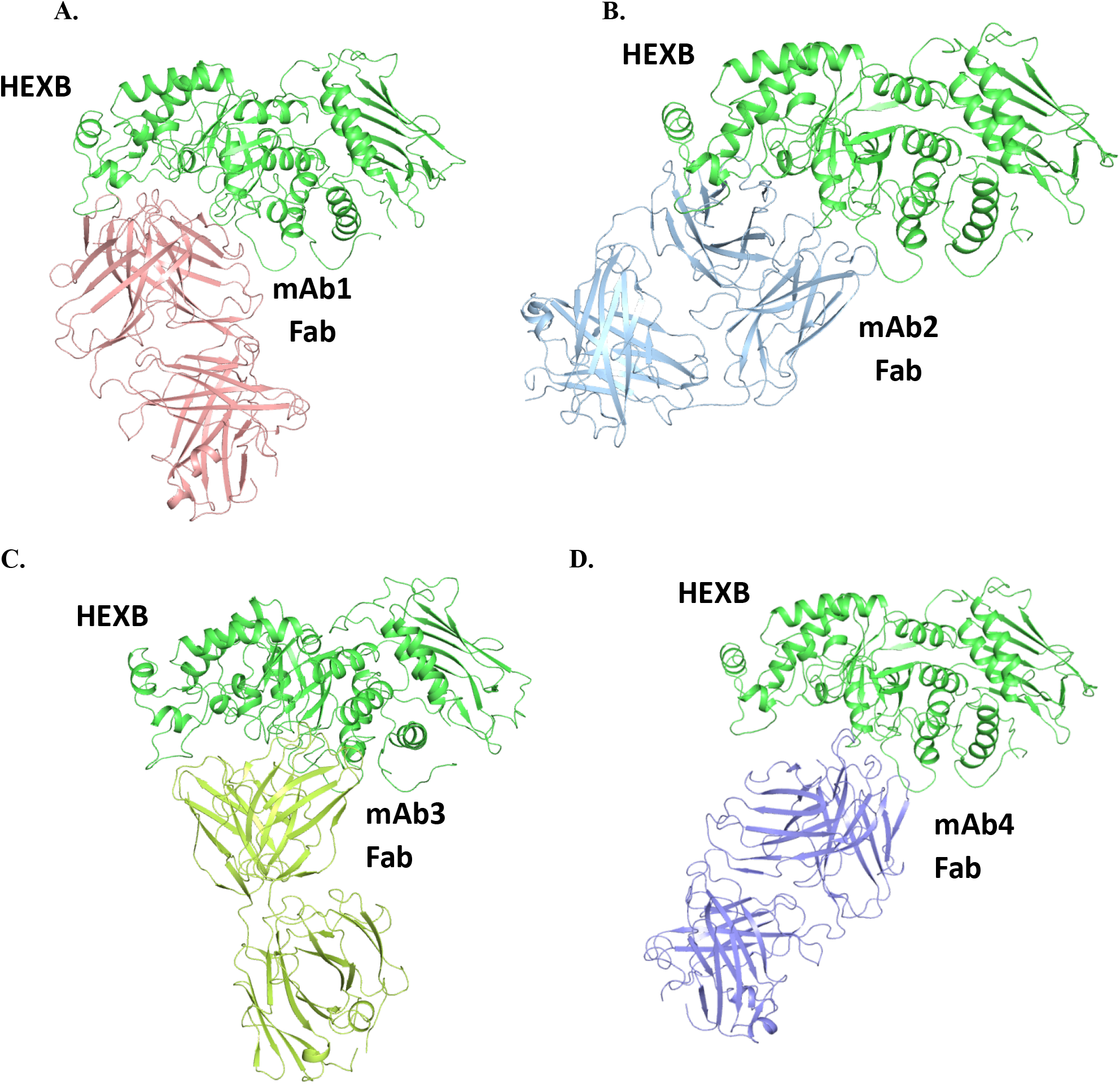
Complex structural models of HEXB bound antibody Fab domains. The complex structures were generated using MOE protein-protein docking with the homology models of HEXB from human and antibody Fab domains based on MOE docking score. While an ensemble of possible interacting pairs is generated by this workflow, only the best scoring model is shown for each pair.

Next, the MMGBSA approach (Molecular Mechanics, Generalized Born model and Solvent Accessibility method) was applied to compute the binding free energy between HEXB and antibodies and to rank the relative binding affinity of all four antibodies. As shown in Table 6, these interaction energy estimates indicate mAb-1 has the lowest binding free energy to HEXB compared to other antibodies, implicating tighter binding of mAb-1 toward HEXB. This is consistent with proteomics observations that HEXB was only detected in mAb-1 DS and causes degradation of N-glycans on mAb-1. Consistent with binding energy, a larger buried surface area is calculated for the HEXB/mAb-1 interface compared to other mAbs, which supports stronger interactions between HEXB and mAb-1.

**Table 6.** Binding affinity between HEXB and antibodies from MMGBSA energy and surface area calculations.

| Complex | Binding Free Energy (kcal/mol) | Buried Surface Area (Å<sup>2</sup>) |
| --- | --- | --- |
| HEXB / mAb-1_Fab | -175 | 1132 |
| HEXB / mAb-2_Fab | -97 | 789 |
| HEXB / mAb-3_Fab | -132 | 649 |
| HEXB / mAb-4_Fab | -146 | 600 |

### Risk Assessment of HEXB in mAb1 on Immunogenicity and Product Quality

To understand the potential immunogenicity risk of HEXB, an *in silico* analysis of the risk posed by the HEXB protein was performed using EpiVax’s ISPRI_HCP algorithm. The relative risk of HEXB was compared to other known HCPs commonly co-purified with mAbs. As shown in Table 7, the EpiMatrix score of HEXB was lower than CXCL3, PLBL2, and HtrA serine peptidase 1. The number of EpiMatrix hits, or 9-mers with a predicted Z-score ≥1.64 that were unique to the CHO protein, and cross-conserved with a human homolog were compared. The HEXB sequence was associated with a higher number of unique hits than PLBL2, a known immunogenic protein, suggesting that it could pose an immunogenicity risk. Anti-PLBL2 antibodies have been detected in individuals dosed with a therapeutic monoclonal antibody that contained different levels of PLBL2.^3^ However, it was noteworthy that the anti-PLBL2 antibodies did not increase the incidence of anti-drug antibodies and did not correlate with safety events.

**Table 7.**
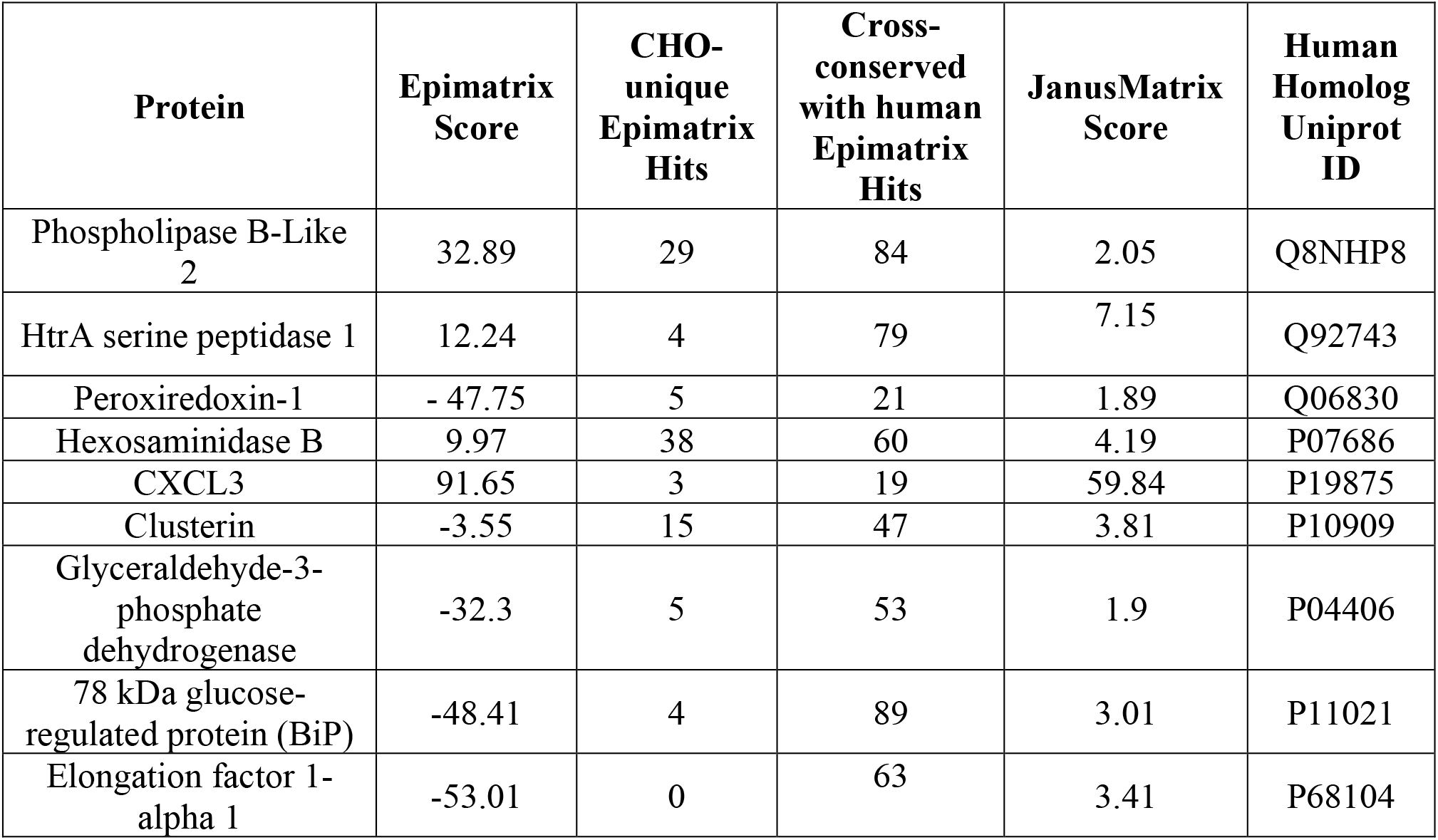
*In silico* analysis of immunogenicity risk of HEXB. Some common problematic HCP was served as controls.

| Protein | Epimatrix Score | CHO-unique Epimatrix Hits | Cross-conserved with human Epimatrix Hits | JanusMatrix Score | Human Homolog Uniprot ID |
| --- | --- | --- | --- | --- | --- |
| Phospholipase B-Like 2 | 32.89 | 29 | 84 | 2.05 | Q8NHP8 |
| HtrA serine peptidase 1 | 12.24 | 4 | 79 | 7.15 | Q92743 |
| Peroxiredoxin-1 | - 47.75 | 5 | 21 | 1.89 | Q06830 |
| Hexosaminidase B | 9.97 | 38 | 60 | 4.19 | P07686 |
| CXCL3 | 91.65 | 3 | 19 | 59.84 | P19875 |
| Clusterin | -3.55 | 15 | 47 | 3.81 | P10909 |
| Glyceraldehyde-3-phosphate dehydrogenase | -32.3 | 5 | 53 | 1.9 | P04406 |
| 78 kDa glucose-regulated protein (BiP) | -48.41 | 4 | 89 | 3.01 | P11021 |
| Elongation factor 1-alpha 1 | -53.01 | 0 | 63 | 3.41 | P68104 |

We next compared the JanusMatrix score of HEXB relative to other known CHO HCPs. This analysis compares the TCR facing amino acids of MHC class II epitopes from the HCPs and predicts cross-reactive epitopes that may be more likely tolerated.^19^ The HEXB sequence was associated with a high Janus score when compared to difficult-to-remove HCPs with the exceptions of CXCL3, Clusterin, and HtrA serine peptidase 1. This score implies that even though the identified epitopes in HEXB could bind with high affinity to prevalent MHC class II molecules, this risk is offset due to the likelihood that CD4+ T cells with TCRs that recognize these cross-reactive epitopes may be clonally deleted or have a regulatory phenotype. Finally, if anti-HEXB antibodies are generated in humans, their risk on patient safety may be low if the target is not easily accessible. For example, the human homolog of CHO PLBL2 is primarily located in the lysosome and may not be recognized by anti-PLBL2 antibodies. This may explain why there were no safety issues linked to the levels of PLBL2 which was related to the incidence of anti-PLBL2 antibodies.^3^ The human homolog of HEXB is reported to be primarily in the lysosome.^20^ Membrane and extracellular forms have been reported as well.^20^ Since both HEXB and PLBL2 are primarily located in the lysosome, the potential generation of anti-HEXB antibodies may have similar safety risks.

The immunogenicity risk of HEXB is deemed to be no more than difficult-to-remove HCPs, but the final assessment will need to evaluate the mAb mechanism of action, disease indication, and take dosing route, amount, and frequency of administration into consideration.

The impact of HEXB on mAb1 product quality is considered low since the N-glycan profile remained stable at the intended storage condition 5 °C. Furthermore, HEXB level is significantly lower and the N-glycan stability is maintained at all stability temperatures after process improvement.

In together, the severity of HEXB in mAb-1 is considered low on potential patient and product risk. Although the occurrence of HEXB in mAb-1 is high as shown in Table 4, we have developed several analytical tools (LC-MS based proteomics and MRM assay, and enzymatical assay) to detect and measure its level and activity. Overall, the risk of HEXB impacting patient safety and product quality related to mAb-1 is deemed low.

## Discussion

This study reported the identification and characterization of a residual HCP, HEXB, leading to degradation of N-glycans on biologics under accelerated storage conditions. The study highlighted the importance of various analytical tools for root cause investigation and control to study residual HCPs in biologics.

HCPs must be adequately removed from biologics by downstream purification to ensure product quality and safety.^1–2^ It is critical to develop phase-appropriate testing for HCP clearance, and the levels are controlled by phase-appropriate specifications. ELISA release assay for HCP testing is relatively simple and high throughput. The assay, however, has several limitations. First, the reported HCP level is highly dependent on the quality and coverage of the antibody reagents used. Additionally, ELISA may entirely miss detecting an abundant HCP if the antibody against it is not present in the critical reagents.^1–2^ Second, the method conventionally used to determine the coverage is a comparison of a two-dimensional (2D) gel to 2D Western blot; both have limitations.^21^ There may be tens or even hundreds of proteins in a single spot in the 2D gel. Thus, the number of spots between gel staining and Western blot may be significantly different. Third, the number of total HCPs measured by ELISA is dilution-dependent and non-linear in some cases. This results from the limited number of antibodies per HCP species. Fourth, because of the specificity of antibody/antigen binding conditions and availability of HCP epitopes, the corresponding HCP may not be readily detected in the ELISA assay even if the correct antibodies are present in the reagent. Lastly, the time to develop a qualified, critical reagent for total HCP measurement by ELISA may take several years. The residual total HCP level by a CHO ELISA kit from Cygnus technologies in the Process 1 of mAb1 is below 30 ppm, which is well below the generally accepted limit (100 ppm).^2^ However, a major change of product quality attribute was observed due to the impact of an abundant residual HCP. This case study, along with many other published reports,^3–8^ emphasized the critical role of orthogonal methods for HCP characterization to ensure the quality of the biologic product.

LC-MS based proteomics approach for HCP characterization is widely used as it is potentially capable of detecting almost any HCP whose abundance is above a certain threshold. The proteomics approach is not only able to identify each HCP, but also provides the relative abundance level of the identified HCP with suitable reference standards. This approach is very informative for the development of customized risk assessment and control strategy for those potential problematic HCPs. In this study, residual HCP HEXB was identified by the proteomics approach as the root cause for N-glycan degradation in mAb-1 from Process 1. The abundance level was determined to be over 200 ppm, which was calculated based on its MS1 peak area compared to the spike-in recombinant non-CHO proteins. Although this approach may suffer accuracy and precision limitations due to the nature of the quantification approach, it provides a quick and fit-for-purpose readout of HCP identification and relative quantification. To compliment the shotgun proteomics approach for quantification, a targeted LC-MS based MRM approach with spike-in stable isotope-labeled internal standards was developed for process and product characterization.^17^ The MRM data provides more accurate measurement of the absolute level of HEXB in mAb1 DS. The HEXB level from MRM assay provides reference for acceptance ranges of HEXB in biologics development. Protein abundance may not correlate with protein activity in a drug product due to buffer conditions or association with other proteins or formulation excipients. Ultimately, it is the enzyme activity which matters the most for its impact on product or excipient degradation. In this study, an enzymatic assay for HEXB was also developed, which provides an alternative assay for mAb-1 process and product characterization. The activity of HEXB from the enzymatic assay in general agrees with the HEXB abundance from both proteomics and MRM assays.

Even after multiple purification steps in the downstream process development, HCPs may co-purify with the final drug product. There is a list of problematic HCPs, which are commonly detected in majority of mAbs by proteomics approach and HEXB is not one of them.^22^ To the best of our knowledge, HEXB is the first residual HCP reported to have impact on the glycan profile of a mAb product in storage conditions. Recently, a co-purifying HCP N-(4)-(β-acetylglucosaminyl)-L-asparaginase (AGA, EC3.5.1.26) was identified by proteomics to explain the unexpected ELISA results. However, no impact on glycan cleavage was reported.^23^ Functional hexosaminidase are dimeric (homodimer or heterodimer) in structure through the combination of α and β subunits (HEXA and HEXB).^20^ Both subunits can cleave GlcNAc residues. Only HEXB was identified in this study. The potential specific interaction of HEXB with mAb-1 leading to co-purification, but not with other mAbs during the downstream purification process was investigated by computational modeling, which provides a structural basis for co-purification and guidance for downstream purification to disrupt the interaction. The potential interactions at the amino acid level will need further validation by other experimental approaches, such as hydrogen-deuterium exchange MS (HDX-MS) or X-ray crystallography. Those validation experiments will heavily rely on the availability of high-qualify recombinant HEXB protein. Furthermore, we are exploring prospective validation of energetic cut-offs, or computational filtering, which would allow for qualitative risk-assessment of HCP interactions needing downstream purification.

In conclusion, a residual HCP, HEXB, was identified to degrade N-glycan by removing terminal GlcNAc from mAb-1 on an accelerated stability study by LC-MS based proteomics approach. The impact was confirmed by enzymatic assay and spike-in recombinant enzyme. The absolute level of HEXB in mAb-1 was determined by LC-MS based MRM assay, and the potential interaction between HEXB and mAb-1 was illustrated by computational modeling. Additionally, the risk assessment of potential impact of HEXB on products and patients were evaluated. This study provided a new case on the impact of residual HCP on biotherapeutics development and highlighted multiple analytical tools for residual HCP characterization and control.

## Acknowledgments

We thank the process team for providing the material and fruitful discussions. We are grateful to Jennifer M. Pollard and Shannon Rivera for reviewing and feedback of the manuscript, and the summer intern Xiujuan (Sophie) Peng for the initial contribution to this work.

---

**Table 1.**
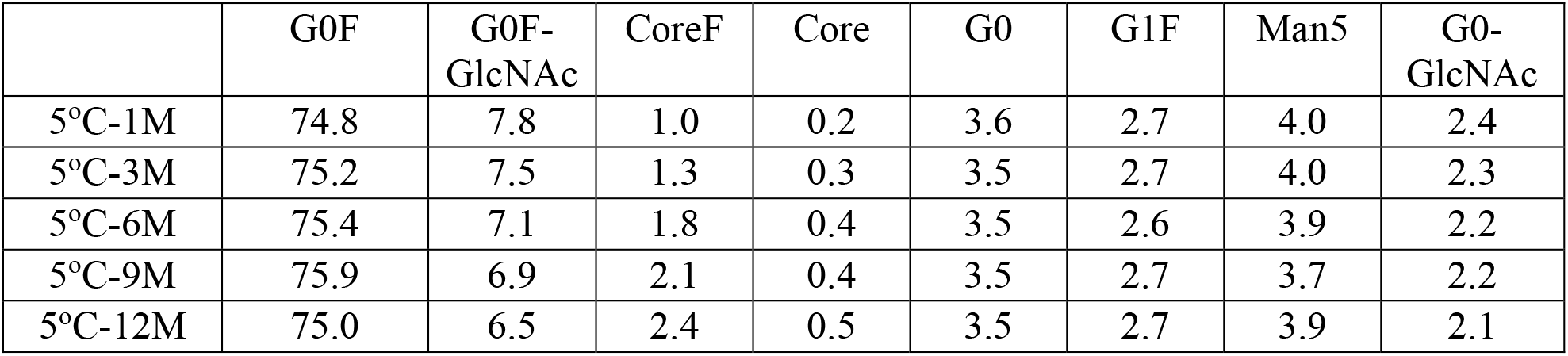
Percent peak area for N-glycan profile of mAb-1 stability sample at 5 °C measured by Instant PC assay.

**Table 2.** Percent peak area for N-glycan profile of mAb-1 stability sample at 25 °C measured by Instant PC assay.

|  | G0F | G0F-GlcNAc | CoreF | Core | G0 | G1F | Man5 | G0-GlcNAc |
| --- | --- | --- | --- | --- | --- | --- | --- | --- |
| 25°C-1M | 74.5 | 6.4 | 3.1 | 0.7 | 3.5 | 2.7 | 4.0 | 1.9 |
| 25°C-3M | 73.0 | 5.4 | 5.7 | 1.4 | 3.5 | 2.6 | 4.1 | 1.2 |
| 25°C-6M | 71.9 | 5.9 | 6.5 | 1.7 | 3.5 | 2.5 | 4.0 | 0.7 |

